# Permutation-based significance analysis reduces the type 1 error rate in bisulfite sequencing data analysis of human umbilical cord blood samples

**DOI:** 10.1101/2021.05.18.444359

**Authors:** Essi Laajala, Viivi Halla-aho, Toni Grönroos, Ubaid Ullah, Mari Vähä-Mäkilä, Mirja Nurmio, Henna Kallionpää, Niina Lietzén, Juha Mykkänen, Omid Rasool, Jorma Toppari, Matej Orešič, Mikael Knip, Riikka Lund, Riitta Lahesmaa, Harri Lähdesmäki

## Abstract

**Background:** DNA methylation patterns are largely established in-utero and might mediate the impacts of in-utero conditions on later health outcomes. Associations between perinatal DNA methylation marks and pregnancy-related variables, such as maternal age and gestational weight gain, have been earlier studied with methylation microarrays, which typically cover less than 2 % of human CpG sites. To detect such associations outside these regions, we chose the bisulfite sequencing approach.

**Methods:** We collected and curated all available clinical data on 200 newborn infants; whose umbilical cord blood samples were analyzed with the reduced representation bisulfite sequencing (RRBS) method. A generalized linear mixed effects model was fit for each high coverage CpG site, followed by spatial and multiple testing adjustment of P values to identify differentially methylated cytosines (DMCs) and regions (DMRs) associated with clinical variables such as maternal age, mode of delivery, and birth weight. Type 1 error rate was then evaluated with a permutation analysis.

**Results:** We discovered a strong inflation of spatially adjusted P values through the permutation analysis, which we then applied for empirical type 1 error control. Based on empirically estimated significance thresholds, very little differential methylation was associated with any of the studied clinical variables, other than sex. With this analysis workflow, the sex-associated differentially methylated regions were highly reproducible across studies, technologies, and statistical models.

**Conclusions:** The inflation of P values was caused by a common method for spatial adjustment and DMR detection, implemented in tools comb-p and RADMeth. With standard significance thresholds, type 1 error rates were high with both these implementations, across alternative parameter settings and analysis strategies. We conclude that comb-p and RADMeth are convenient methods for the detection of differentially methylated regions, but the statistical significance should either be determined empirically or before the spatial adjustment. Our RRBS data analysis workflow is available in https://github.com/EssiLaajala/RRBS_workflow.

## Background

Mitotically inheritable DNA methylation patterns are established in early embryogenesis and can be influenced by environmental and lifestyle-related factors (1). The in-utero environment might be the most important explanatory factor for the between-individual variation in genome-wide average DNA methylation (2). Several human and animal studies have identified DNA methylation to mediate the impacts of in-utero conditions on later health (3,4). Correlations between maternal and pregnancy-related factors and perinatal DNA methylation have been actively studied during the past decade. Associations of umbilical cord blood DNA methylation marks with maternal smoking, maternal BMI, birth weight, gestational age, and maternal gestational diabetes have been mapped in large meta-analyses of methylation microarray data sets from multiple study cohorts (5–9). All these factors were associated with some differential CpG methylation. Maternal smoking during pregnancy was associated with thousands of differentially methylated cytosines, all of which showed some evidence of differential methylation in older children as well, indicating that DNA methylation patterns are relatively stable with respect to age (5). Most earlier studies on umbilical cord blood DNA methylation have been limited to a set of approximately 450 000 CpG sites covered by the methylation microarrays.

We applied reduced representation bisulfite sequencing (RRBS) on 200 umbilical cord blood samples to accomplish a genome-wide survey on associations between perinatal DNA methylation and various clinical covariates. These included e.g. maternal age, gestational weight gain, mode of delivery, and the birth weight of the newborn infant (full list in Table 1, details in Supplementary Table 1). Initially, applying a standard analysis workflow, hundreds of differentially methylated CpG sites were associated with each of these clinical variables. As a plausibility test, the analysis was repeated for a permuted variable, which did not correlate with any clinical or technical variable and should ideally not be associated with any differential methylation. A strong inflation of spatially adjusted P values was observed for the permuted variable. Importantly, targeted validation by pyrosequencing confirmed the lack of significant differential methylation in candidate regions that were selected based on standard criteria (Benjamini-Hochberg corrected spatially adjusted P value < 0.05) before the P value inflation was discovered (10). Here, we further explore this phenomenon with different study designs and analysis workflows, and present practical recommendations, including implementations as R code, for future bisulfite sequencing studies. We report appropriately FDR (false discovery rate) controlled results on associations between perinatal DNA methylation and the above-listed variables within and beyond the genomic locations covered by earlier studies.

**Table 1:** Numbers of differentially methylated cytosines and regions associated with each covariate. Column 1 contains the experimentally determined threshold values for spatially adjusted P values and column 2 contains the corresponding numbers of CpG sites within differentially methylated regions (DMRs). The threshold is set such that the number of findings associated with a permuted version of the covariate would be less than 5 % of the number of findings associated with the original covariate (column 2). The DMRs in column 3 are defined as described in Methods, filtering for the concordance of the direction of difference. Column 5 contains the numbers of differentially methylated cytosines (DMCs) detected without spatial adjustment (Benjamini-Hochberg-corrected PQLseq P value < 0.05), some of which belong to differentially methylated regions, and column 4 contains a subset of the DMCs in column 5.

|  | Adjusted P value threshold (median of 3 permutations) | Number of CpG sites within DMRs (spatial adjustment + empirical FDR control) | Number of DMRs | Number of DMCs outside DMRs (FDR < 0.05 before spatial adjustment) | Total number of DMCs (FDR < 0.05 before spatial adjustment) |
| --- | --- | --- | --- | --- | --- |
| Year of birth (= sample collection year) | 0 | 0 | 0 | 6 | 6 |
| Smoking during pregnancy, mother, N=14 | 0 | 0 | 0 | 1 | 1 |
| Sex, N=68 (females) | 4.67E-07 | 6165 | 297 | 261 | 1426 |
| Month of birth (cosine transformed) | 0 | 0 | 0 | 0 | 0 |
| Insulin-treated diabetes, mother, N=8 | 1.13E-10 | 29 | 2 | 10 | 10 |
| Induced labor, N=28 | 0 | 0 | 0 | 0 | 0 |
| Height, mother | 6.55E-11 | 47 | 2 | 3 | 3 |
| Gestational weight gain | 9.53E-13 | 16 | 1 | 0 | 0 |
| Epidural anesthetic, delivery phase 1, N=86 | 1.22E-15 | 3 | 1 | 0 | 0 |
| Earlier miscarriage(s), N=31 | 0 | 0 | 0 | 0 | 0 |
| Caesarean section, N=21 | 0 | 0 | 0 | 0 | 0 |
| BMI, mother (pre-pregnancy) | 0 | 0 | 0 | 0 | 0 |
| Birth weight | 0 | 0 | 0 | 0 | 0 |
| Apgar points low, 1 minute, N=24 | 3.99E-12 | 19 | 2 | 2 | 2 |
| Age, mother | 9.59E-12 | 29 | 2 | 0 | 0 |

Accounting for spatial correlation between CpG sites is important in all DNA methylation studies and especially in the analysis of bisulfite sequencing data, where methylation is quantified on single-nucleotide resolution. One of the simplest approaches is to first compute P values for differential methylation at each CpG site separately and then to combine them with an autocorrelation-adjusted Z-test, also known as the Stouffer-Lipták-Kechris correction, implemented in the Python package comb-p (11). A Stouffer-Lipták meta-P-value is a (anti-probit-transformed) weighted sum of probit-transformed P values, developed for meta-analyses of results from multiple independent studies (12,13). Kechris et al. suggested the application of a similar approach to combine adjacent P values in spatially correlated genomic data, more specifically in tiling array data (14). Since such P values do not meet the independence assumption of the Stouffer-Lipták method, Kechris et al. suggested adjustment for autocorrelation. Comb-p is a generalization of the Stouffer-Lipták-Kechris method to non-evenly spaced spatially correlated P values (11). It has become a popular tool especially for the detection of differentially methylated regions (DMRs) in DNA methylation microarray studies. It is also part of the bisulfite sequencing data analysis pipeline MethPipe (15), as implemented by the developers of the beta-binomial regression model RADMeth (16).

Comb-p (11) estimates the autocorrelation between P values up to a chosen genomic distance (for example 500 bp), performs a sliding window Stouffer-Lipták-Kechris correction for each P value, utilizes a peak detection method to detect potential DMRs, and assigns P values to these variable-sized regions. The region-wise P values are calculated by repeating the Stouffer-Lipták-Kechris correction for each candidate region, after which they are corrected for multiple testing with a Sidák-correction, based on the number of times the largest candidate region would fit in the total number of features (such as CpG sites). The spatial adjustment method within RADMeth (16) includes the same autocorrelation estimation step and the Stouffer-Lipták-Kechris correction with a sliding window, recommended to be of size 200 bp. RADMeth performs a Benjamini-Hochberg-correction for the spatially adjusted P values and detects regions by finding consecutive CpG sites with FDR < 0.01. The specificity of this DMR detection method has been earlier evaluated in simulated data but not in real bisulfite sequencing data with a permutation analysis, as presented here.

## Methods

The methods are summarized in Figure 1 and have been partially described in a preprint by Laajala et al. (10). Most of the data analysis and visualization were done with R versions 3.6.3 and 4.0.4. (17). The R code and a documentation of the usage of command-line tools are available in GitHub (18). R packages Hmisc (19) and gplots (20) were used to generate Figure 2, ggplot2 (21) was used to generate Figure 3, and stringr (22) was used for some basic string processing.

**Figure 1:**
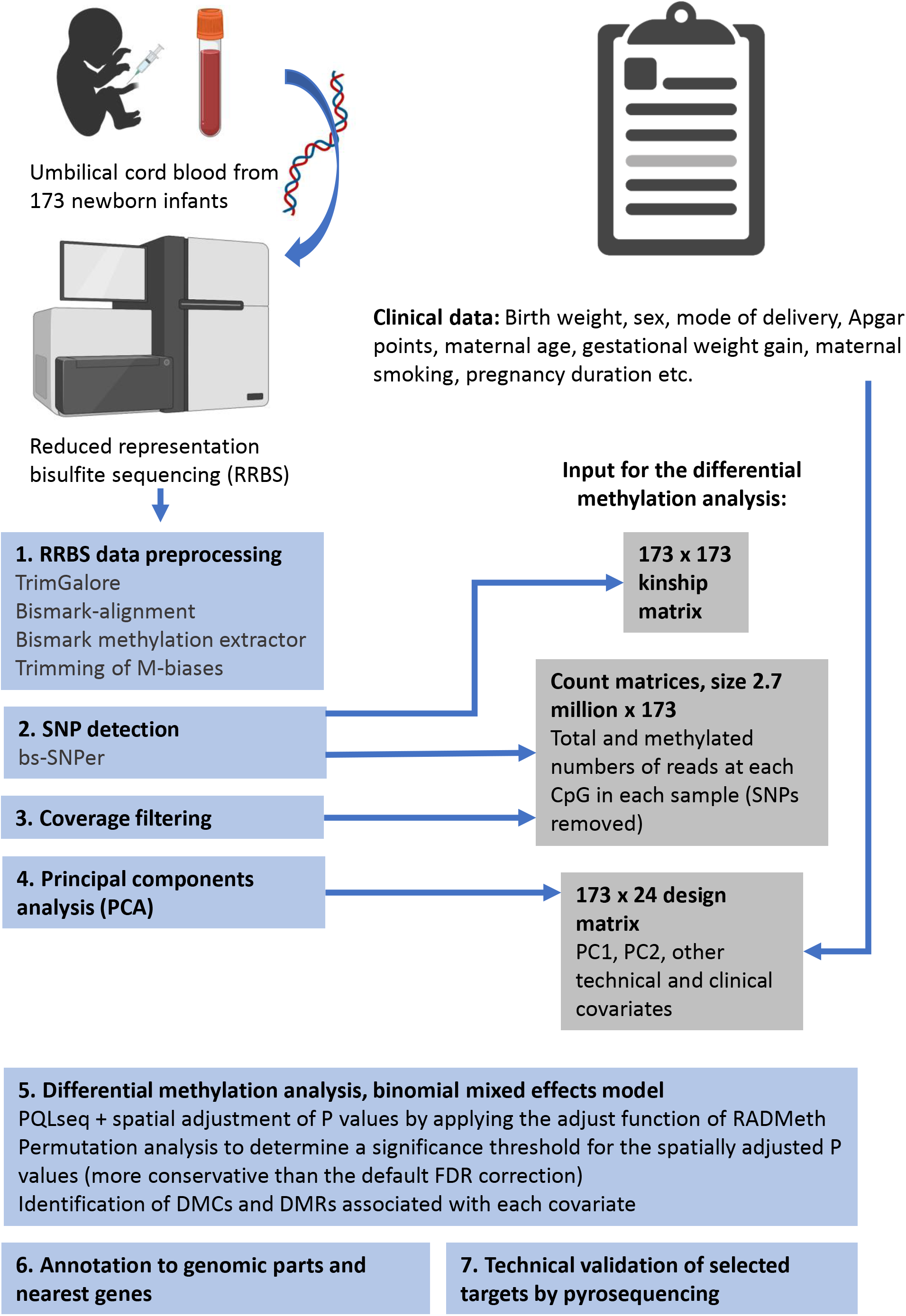
Outline of the study design and methods, created with BioRender.com

**Figure 2:**
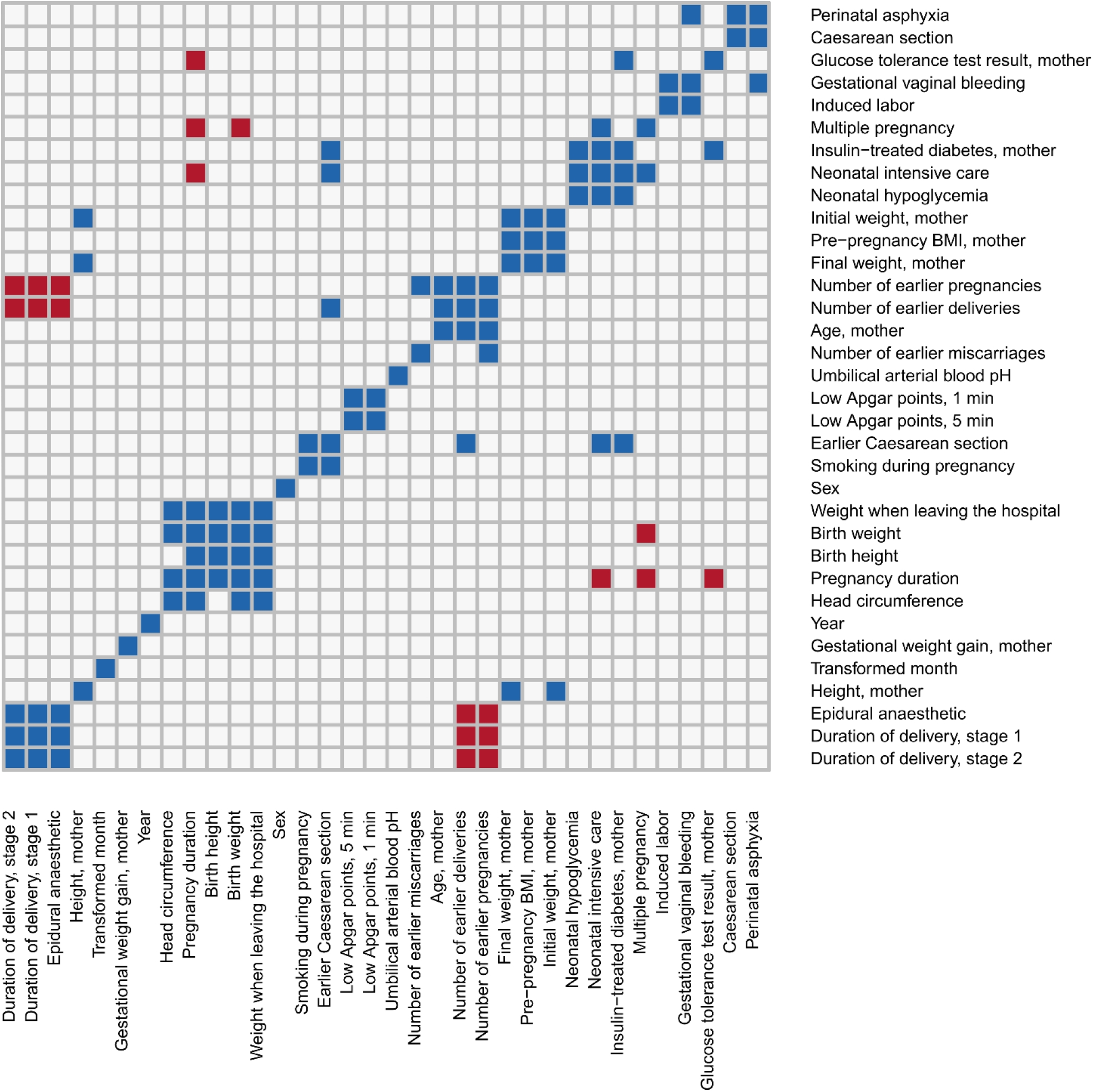
Observed correlations between clinical covariates in these data. Blue and red color indicate significant (P value < 0.05, absolute Pearson’s r > 0.3) positive and negative correlation, respectively. For each pair of binary covariates, significance was determined using Fisher’s exact test (P value < 0.05). Binary/categorical variables with 5 or fewer examples of a category are excluded from this figure.

**Figure 3:**
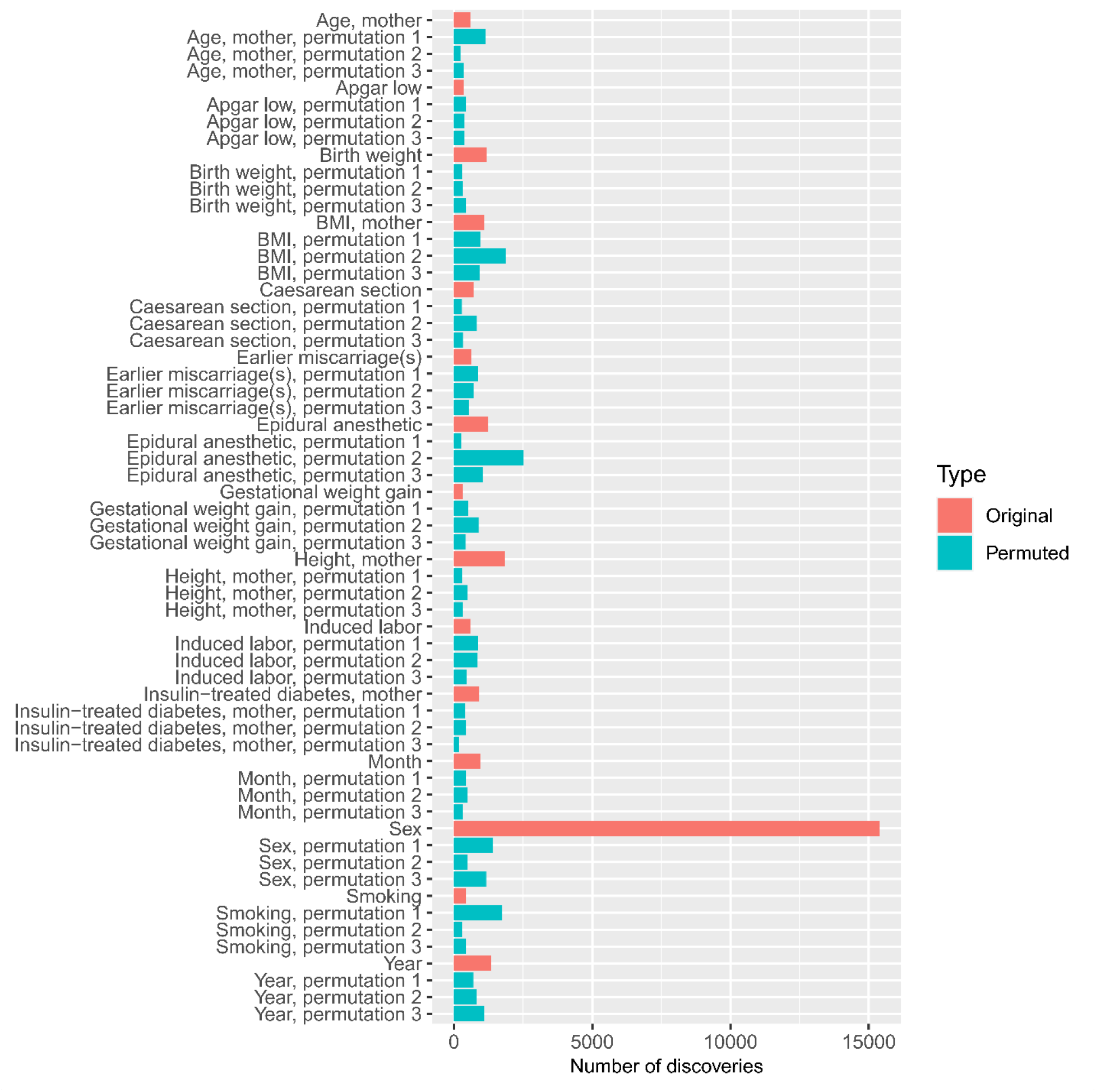
The numbers of differentially methylated CpG sites associated with each original and permuted covariate would have been these, if the default significance threshold (Benjamini-Hochberg corrected spatially adjusted P value < 0.05) had been applied. These numbers were obtained by performing a differential methylation analysis (fitting a GLMM to obtain a Wald test P value for each CpG site, followed by the spatial adjustment and multiple testing correction implemented within RADMeth) for the original input data, as well as for 45 permuted design matrices (3 permutations of each of the 15 covariates of interest). This permutation analysis showed that the spatially adjusted P values were inflated.

### Umbilical cord blood DNA collection and reduced representation bisulfite sequencing (RRBS)

The genome-wide umbilical cord blood DNA methylation measurements are from a study by Laajala et al. (10), which includes a detailed description of the study design, sample collection, and reduced representation bisulfite sequencing (RRBS). Briefly, these data were collected from participants of the Finnish Diabetes Prediction and Prevention (DIPP) follow-up study, who were born in Turku University Hospital between 1995 and 2006. The umbilical cord blood samples were collected immediately after birth in 3 ml K3-EDTA tubes, transferred to the DIPP clinic, and stored at −20°C. DNA was extracted by the salting out procedure (23) and purified with Genomic DNA Clean & Concentrator kit (Zymo Research, cat. nos D4010 and D4066) according to the manufacturer’s protocol. RRBS library preparation steps were adapted from (26,27). Aliquots of bisulfite converted DNA were amplified by 18 cycles of PCR and sequenced with Illumina HiSeq 2500 instrument using paired-end sequencing with read length 2 × 100 bp. The technical quality of the HiSeq 2500 run was good and the cluster amount was as expected. The yields were 18 - 37 million raw paired-end reads per sample. Out of 200 umbilical cord blood DNA samples, five were rejected due to inadequate amount or quality of DNA, 20 were later rejected due to low (< 97 %) bisulfite conversion efficiency, and two were excluded due to missing clinical data.

### Clinical data collection and curation

All available data on pregnancy and birth were retrieved from Turku University Hospital. This included variables related to the mother (such as age and number of earlier pregnancies), the pregnancy (such as glucose tolerance test result and gestational weight gain), the delivery (such as perinatal asphyxia and usage of epidural anesthetic), and the child (such as birth weight and neonatal intensive care). The available clinical covariates are listed in Supplementary Table 1. All technical covariates were included in the differential methylation analysis (described below), and clinical covariates were selected such that within each group of mutually correlating covariates, the most reliably measured covariate was included. For example, birth weight was selected to represent the size of the newborn infant, while birth length and head circumference were excluded. Pearson correlations greater than 0.3 (in absolute value) with P value < 0.05 or alternatively Fisher’s exact test P value < 0.05 (for pairs of binary covariates) were considered relevant. Further details are in Supplementary Table 1 and Figure 2.

The clinical data required some manual curation. A cosine transformation cos(2πm/12) was applied to account for the cyclic nature of the month of birth (m = month). Apgar points were simplified to normal/low such that values 8 - 10 were considered normal. Mode of delivery was originally a multi-level categorical variable with different values for normal, forceps, suction cup, elective Caesarean section (C-section), urgent C-section, and emergency C-section but was simplified to C-section/vaginal. Since the data only included two women who smoked only during the first trimester and 12 women who smoked throughout the pregnancy, this variable was simplified to smoking/no smoking. For similar reasons, the number of earlier miscarriages and the number of fetuses were simplified to binary variables (twins and triplets were all marked as multiple pregnancies). Usage of epidural anesthetic during delivery phase 1 was corrected to 0 for two individuals with an elective C-section.

For the regression modeling, missing covariate values were median-imputed, which was necessary only for smoking during pregnancy (four missing values), gestational weight gain (three missing values), maternal pre-pregnancy BMI (two missing values), and Apgar points (one missing value). Variable selection and the visualization in Figure 2 were carried out before any imputation of missing values but after the above-mentioned simplifications and corrections. Continuous covariates were Z-transformed (divided by the standard deviation after subtracting the mean) for the regression model.

### Read trimming and sequence alignment

The sequencing reads were trimmed using TrimGalore version 0.4.3 (25) in paired-end RRBS mode, which removes end repair biases by default. Quality control was done by examining fastQC reports generated before and after running TrimGalore. According to these reports, TrimGalore had correctly removed end repair biases, adapters (adapter sequence minimum overlap 1), and bases with base call error rate above 1%. Sequence duplication levels were elevated, as expected in the context of RRBS, but adapters were not among the overrepresented sequences after running TrimGalore. By default, TrimGalore discards reads shorter than 20 bp after trimming. This step removed 2 - 8 % of the raw reads from all samples except one sample, from which 24 % of raw reads were discarded.

The reads were aligned on the human GRCh37 (hg19) genome assembly (26,27) and the lambda phage genome simultaneously with Bowtie2 version 2.3.1 within Bismark version 0.17.0 (28) after preparing the genome using function Bismark_Genome_Preparation, which creates bisulfite converted versions of the genome (both C - > T and G - > A converted versions of each genomic area). These steps were run using the default parameters of Bismark in paired-end mode. The documentation of this project in GitHub (29) includes the precise commands and parameters.

### Methylation call extraction, conversion efficiency calculation, and removal of M-biases

To extract the number of methylated and unmethylated reads at each CpG site in each sample, Bismark methylation extractor version 0.22.3 (28) was run with parameters paired-end, bedGraph, and counts. To avoid redundant methylation calls within pairs of reads, this function (by default) excludes read 2 bases that overlap with read 1. The extracted counts within the lambda phage genome were used to determine the conversion efficiency of each sample (the sum of observed unmethylated CpG counts divided by the total sum of methylated and unmethylated CpG counts within this completely unmethylated genome). The conversion efficiencies were above 97 % (median 99.4 %) for all except 20 samples, which were therefore excluded from the analyses.

Bismark methylation extractor outputs M-bias-files, which contain context-specific (CpG, CHG, and CHH) average methylation proportions in each read position. To remove technical methylation call artefacts in the ends of sequencing reads, we used the middle positions to determine the normal variation of CpG-specific methylation and removed anything that was beyond that, as suggested by the authors of BSeQC (30). Specifically, positions at read ends were removed if their CpG methylation proportions were more than 3 standard deviations below or above the mean methylation proportion at positions 10 – 91 (middle 80 %). However, the 5’ ends of read 1 were not trimmed, since they start with a CpG site and typically have higher average methylation levels than other positions. This is explained by the fact that a CpG site is always present at the beginning of an RRBS read even if the read comes from a CpG-poor area, whereas CpG sites at middle positions are more likely to come from CpG-rich areas, which typically have lower methylation levels. In these data, three bases were trimmed from the 5’ end of read 2 and one base from the 3’ end of read 2 due to CpG-specific M-biases in every sample. This was done by re-running the Bismark methylation extractor as above with additional parameters ignore_r2 3 and ignore_3prime_r2 1. Finally, the information from both strands was merged for each CpG site by running Bismark function coverage2cytosine with parameter merge_CpG for each cov-file produced by the Bismark methylation extractor.

Ideally CHH- and CHG-specific M-bias profiles should be flat lines close to zero. The only samples with CHH-or CHG-specific methylation above 1 % (at any position) had already been excluded when the lambda phage genome was used to determine the bisulfite conversion efficiency. Therefore, no further exclusions were done at this step.

### Count matrix construction and SNP removal

The numbers of methylated and unmethylated reads were extracted from merged_CpG_evidence.cov files produced by Bismark function coverage2cytosine (28) and organized into count matrices with total and methylated numbers of reads for each CpG site and each sample. To complete this step within a reasonable time and memory reservation, only CpG sites with minimum coverage of 10 in at least ten samples were included. However, the actual coverage filtering was done after SNP removal and is described below. The dimensions of the pre-filtered count matrices created at this stage were 3 928 420 CpG sites × 173 samples.

SNP detection was done by applying bsSNPer (30) with its default parameters on bam-files sorted by genomic coordinates after excluding the lambda phage genome. The detected SNPs (flagged “PASS” in the bsSNPer output VCF file) were removed from the data, specific to each individual (read counts set to NA).

### Coverage filtering

In the context of next-generation sequencing in general, a common practice is to remove PCR duplicates by excluding reads that align at exactly the same genomic coordinates. However, this is not possible for RRBS data, where identical fragments are more likely to originate from different molecules, which were cut at exactly the same positions by MspI. To remove most PCR duplication biases, CpG sites with coverage above the 99.9^th^ percentile were removed from each sample, as suggested by the authors of MethylKit (31).

CpG sites were completely excluded if they had a low-coverage value (total number of reads < 10) or a missing value (a potential SNP or the coverage above the 99.9^th^ percentile of the sample) in at least two thirds of the samples. The below-described differential methylation analysis was run for all 2 752 981 CpG sites passing these criteria. However, in case of binary covariates, further covariate-specific filtering was done before spatially adjusting and FDR-correcting the P values. Minimum coverage of 10 in at least one third of the samples in each group was required and further, a minimum coverage of 10 was required in at least five samples per group (the second criterion is relevant only for binary covariates with less than 15 samples in one group).

### Principal components analysis (PCA)

To be able to perform principal components analysis (PCA) on methylation proportions (methylated/total reads), missing values at each CpG site were imputed by the median over samples with non-missing values. After removing CpG sites in chromosomes X and Y, we applied a readily available implementation of PCA (R-package calibrate version 1.7.5, (32)) with its default parameters on the coverage-filtered imputed methylation proportion matrix. Principal coordinates 1 and 2, i.e. projections of the sample-specific methylation proportion vectors on the first two orthonormal principal components were included as covariates in the differential methylation analysis to correct for technical variation between the samples.

### Detection of differentially methylated CpG sites (DMCs) associated with different covariates

The differential methylation analysis was carried out by applying a generalized linear mixed effects model (GLMM) implemented in R package PQLseq (33) separately for each CpG site. PQLseq models the technical sampling variation in bisulfite sequencing data with a binomial distribution, effects of biological and technical covariates with the linear model, and the random effects with a correlated multivariate normal distribution. To model these data, the main improvement compared to simpler beta-binomial models is that it can include continuous covariates.

PQLseq version 1.1. was applied with R version 3.6.1 on the coverage filtered count matrices (numbers of methylated and total reads of each sample in each CpG site), including only chromosomes 1 - 22. This was done after adding + 1 to the numbers of methylated reads and +2 to the total numbers of reads to avoid modeling methylation proportions that are exactly 0 or 1, as recommended by the authors of PQLseq (33). This pseudo-count transformation was only applied to non-missing values (coverage > 0). The clinical covariates listed in Table 1, the case/control-status from (10) (positivity for islet autoantibodies before age 15), library preparation batch, and principal components 1 and 2 were included in the model as fixed effects (detailed descriptions and inclusion criteria of the covariates are presented in Supplementary Table 1). At each CpG site, binary covariates were included only if at least 3 samples with enough coverage (pseudo-count transformed coverage ≥ 12) were available for each category. Including covariates with no data would have unnecessarily caused convergence failures. Since we were interested in differential methylation associated with each covariate (not only a single covariate as assumed in PQLseq implementation), the source code of PQLseq was modified to output the coefficients, standard errors and Wald test P values for all covariates included in the model (the modified version is included in GitHub (18)).

Since PQLseq was originally designed to model differential methylation in the presence of population structures, we included the relatedness of the individuals as a random effect. To our knowledge, these 173 individuals are unrelated, but we estimated their genetic similarity by utilizing the SNPs detected as described above. The relatedness matrix is a correlation matrix of the samples’ SNP profiles, which include all detected (flagged “PASS”) SNPs with minor allele frequency > 5%, encoded as the number of reference alleles (0,1,2). This is calculated as XX^T^/N_SNPs_, where X is a N_samples_ × N_SNPs_ matrix (173 × 187569) containing numbers of reference alleles, standardized to Z-scores within each sample. The number of reference alleles was assumed to be 2, unless a SNP was detected.

### Spatial adjustment and FDR-correction of raw P values

The Wald test P values computed within PQLseq were spatially adjusted by utilizing the adjust-function implemented by the developers of RADMeth (16) within Methpipe version 3.4.3. (15) after sorting the CpG sites by chromosome and location. The recommended window size 200 was used with step size 1. This function performs a Stouffer-Lipták-Kechris adjustment (autocorrelation-adjusted Z-test) of the P values, followed by a Benjamini-Hochberg-correction, but we found this procedure to be insufficient for FDR control. Therefore, the differential methylation analysis (PQLseq) and the spatial adjustment of P values were repeated for permuted input covariates to estimate the null distributions of spatially adjusted P values associated with different types of covariates. For this purpose, 45 (3 × the number of covariates of interest) different input design matrices were created, each with one covariate permuted such that it did not correlate with any actual clinical or technical covariate. Low correlations (absolute Pearson correlation coefficient < 0.3) with continuous clinical covariates were allowed, other than that any significant correlation (P value < 0.05) was considered too strong (permutations were repeated until none was observed). Each distribution of spatially adjusted P values associated with a permuted covariate was compared to that of the corresponding original covariate. A threshold for the spatially adjusted P value was set such that the number of discoveries in permuted data (false discoveries) would be less than 5 % of the number of CpG sites associated with the original covariate using that threshold. The median threshold value over three repeats was used.

CpG sites with the spatially adjusted FDR < 0.05 (using the above-described empirical FDR control) or FDR < 0.05 (Benjamini-Hochberg-corrected PQLseq P value before any spatial adjustment) were included in the DMR analysis described below. CpG sites of the latter type (FDR < 0.05) were reported as differentially methylated cytosines (DMCs) even if they were not part of any DMR.

### Detection of differentially methylated genomic regions (DMRs)

A differentially methylated region (DMR) was defined as a genomic region with two or more CpG sites with evidence of differential methylation (empirically FDR-corrected spatially adjusted P value < 0.05 or Benjamini-Hochberg-corrected PQLseq P value < 0.05) that were within a window of 2 kb and had the same direction of methylation difference in at least 90% of the CpG sites. Table 1 and Supplementary Tables 2 and 3 only include DMRs that had absolute coverage-corrected mean methylation difference > 5 % in at least one CpG site. Coverage-corrected mean methylation difference for a single CpG is calculated as sum(number of methylated reads in cases)/sum(number of total reads in cases) – sum(number of methylated reads in controls)/sum(number of total reads in controls).

### Alternative differential methylation analysis workflows: RADMeth beta-binomial regression and DMR detection with RADMeth and comb-p

For comparison purposes, Type 1 error rate was evaluated with alternative analysis workflows in addition to the one described above. The GLMM for differential methylation at each CpG site (PQLseq) was replaced with a beta-binomial regression model RADMeth (16), implemented as function “radmeth regression” within MethPipe version 3.4.3. Since RADMeth is unable to include continuous covariates, each continuous covariate was represented as two binary covariates: “high” corresponding to the highest quantile and “low” corresponding to the lowest quantile (values in the middle two quantiles were included in the intercept). The count matrices (methylated and total) were pseudo-count transformed, similarly as for PQLseq, and combined to the input format for RADMeth. RADMeth and PQLseq runs were repeated without the pseudo-count transformation and with alternative (simpler) study designs: sex + PC1 + PC2 or epidural + sex + PC1 + PC2. Epidural (the usage of epidural anesthetic during delivery phase I) is an example of a binary covariate with 50 % of the samples in each category that is not expected to be associated with differential methylation (but if differences were present, sample numbers 86 vs. 87 would probably be enough to detect them). CpG sites for which RADMeth did not converge (P value “−1”) were removed from the output. With the pseudo-count transformation, however, the convergence was 100 %. The output was sorted by chromosome and location before the spatial adjustment.

As alternative DMR detection methods, we also used “radmeth merge” (within MethPipe 3.4.3.) with parameter −p 0.01 (to combine adjacent CpG sites with Benjamini-Hochberg-corrected spatially adjusted P values < 0.01 into regions) and the comb-p pipeline (26). The Python package for comb-p was cloned from GitHub (34) (version 0.50.4. accessed on March 25^th^, 2021) and used with Python version 3.7.3. Window size 200 and step size 1 were used for both implementations of the Stouffer-Lipták-Kechris adjustment (also window size 500 was tried with step size 10 but the results were almost identical to those obtained with window size 200). The DMR detection steps of comb-p were performed with seed 0.1 (P value threshold to start a candidate region), and Sidák-corrected region-wise P value < 0.05 (computed by comb-p) was used as a criterion to define a DMR.

### Annotation and gene ontology enrichment analysis of differentially methylated genomic regions

Differentially methylated CpG sites were annotated to genomic parts (promoter, intron, exon, intergenic) and nearest UCSC known genes through R package genomation version 1.16.0 (35) using Genome Reference Consortium Human Build 37 (GRCh37, hg19 (26)). A Gene Ontology enrichment analysis was performed on the list of all nearest genes annotated to DMRs and DMCs associated with sex using the 2020-10-09 Gene Ontology release through PANTHER (36). Significantly enriched biological process gene ontologies (Fisher’s exact test FDR < 0.05) are listed in Supplementary Table 4.

### Pyrosequencing validation of a selected target

Targeted pyrosequencing was performed for technical validation of a DMR on the promoter of zona pellucida binding protein 2 (*ZPBP2*), which is associated with sex. For this analysis, 25 female and 34 male individuals were chosen with the following criteria: full-term (gestational age ≥ 37 weeks), normal birth weight (2.5 – 4.5 kg), no multiple pregnancies, normal Apgar points (8 – 10), no perinatal asphyxia, vaginal birth and no maternal smoking. The sequencing was done on five batches such that a roughly even number of males and females were allocated on each batch. PyroMark assay design 2.0 software (Qiagen) was used to design an assay for the region of interest (chr17:38024237 - chr17:38024291 on hg19 coordinates, including six differentially methylated CpG sites). Sample preparations were started with 200 ng of DNA from the selected samples. Samples were sodium bisulfite treated with EZ DNA Methylation-GoldTM Kit (Zymo Research cat no D5006), and the target sequence was amplified by PyroMark PCR Kit (Qiagen cat no 978703) (initial denaturation at 95 °C for 15 min, 45 cycles at 94 °C for 30 s, annealing at 56 °C for 30 s, extension at 72 °C for 30 s and a final extension at 72 °C for 10 min). Pyrosequencing was carried out on PyroMark Q24 system (Qiagen) using PyroMark Q24^®^ Advanced CpG Reagents (Qiagen cat no 970922), and methylation percentages were extracted from the light intensity values at each CpG site using PyroMark Q24 Advanced software 3.0.1.

### Pyrosequencing data analysis

A standard linear model was applied after transforming each methylation proportion with: arcsin(2×proportion–1). In addition to the sequencing batch, covariates specified in Supplementary Table 1 were included in the model, excluding those that were not applicable to the set of pyrosequenced samples (such as maternal smoking, which was 0 for all these samples). The model was fit with and without sex, and the models were compared with a likelihood-ratio test to assess the significance of sex at each CpG site. These steps were done using functions lm and anova, R version 4.0.4. (17)

## Results

### Observed correlations between clinical and technical covariates

Since fitting a regression model for mutually correlating covariates would be problematic, we selected only one covariate from each group of strongly correlated clinical covariates (Figure 2). The reasons for including and excluding covariates are detailed in Supplementary Table 1. The study was originally not designed for associations between DNA methylation and these clinical variables (instead, the samples were allocated to library preparation batches for optimal comparison between the study groups described in the study by Laajala et al. (10)). However, none of the studied clinical covariates (listed in Table 1) correlated with the library preparation batches or other technical covariates. Technical covariates only correlated with other technical covariates, including the sample collection year.

### Stouffer-Lipták-Kechris-corrected P values were inflated

As a plausibility test, we repeated the differential methylation analysis several times using permuted covariates that did not correlate with any real clinical or technical covariate. The distributions of raw Wald test P values from the binomial mixed effects model PQLseq (33) were as expected: when a permuted input covariate was used, the number of findings (Benjamini-Hochberg-corrected P values below 0.05) was zero. This was not the case for the spatially adjusted P values, which were obtained by applying the adjust-function within RADMeth (16). The number of CpG sites that were considered differentially methylated in each shuffled analysis (false discoveries) was between 200 and 2500 (Figure 3), often close to the number that would have been associated with the corresponding original covariate, if a standard cutoff (Benjamini-Hochberg-corrected spatially adjusted P value < 0.05) had been applied. The distributions of spatially adjusted P values obtained by running each differential methylation analysis with permuted input covariates were utilized to find suitable P value thresholds (see Methods for details). This empirical type 1 error control decreased the number of findings associated with each covariate to zero or a small fraction of the number that would have been discovered with the default FDR control of RADMeth (Table 1 compared to Figure 3).

Since the spatial adjustment that we used was originally proposed to be used together with beta-binomial regression, we repeated a subset of the permutation analysis with the RADMeth workflow. Table 2 contains the numbers of false discoveries, that were observed when the spatial adjustment and DMR detection of RADMeth were applied to P values from the beta-binomial regression model of RADMeth, as compared to P values from PQLseq. These numbers suggest that the RADMeth workflow has an even greater type 1 error than the above-described combination of PQLseq and RADMeth’s spatial adjustment. For example, 3115 – 10555 CpG sites were associated with permuted sex, when the RADMeth workflow was applied (as compared to 482 – 1414, when the spatial adjustment of RADMeth was applied on P values from PQLseq, as shown in Figure 3 and repeated in Table 2). In fact, the beta-binomial regression P values were already inflated before the spatial adjustment. We observed 80 – 813 CpG sites that were associated with permuted sex at Benjamini-Hochberg-corrected P value < 0.05. This was largely explained by inappropriate handling of missing values (coverage = 0) by RADMeth. When coverage was zero at all samples with a given level of a (binary) covariate, RADMeth reported an extremely small P value, when it should have either ignored the covariate, reported a convergence failure, or thrown an error/warning (it does report a failure if this is the case for the covariate of interest but apparently does not check it for the other covariates). In contrast, PQLseq ignores samples with coverage 0 at each CpG site and reports a convergence failure, in case no data remain for some category (we avoided these events altogether by using a CpG-specific design matrix, as described in the Methods). For a fair comparison between RADMeth and PQLseq, we repeated the permutation analysis for simple study designs: permuted sex + PC1 + PC2 and permuted epidural + sex + PC1 + PC2. This almost completely removed the inflation of beta-binomial regression P values and decreased the differences between PQLseq and RADMeth (Table 2 and Table 3). To summarize these results, the observed inflation of spatially adjusted P values was due to the spatial adjustment and had nothing to do with PQLseq.

**Table 2:** Numbers of DMCs and DMRs associated with permuted covariates (false discoveries), when RADMeth’s spatial adjustment and default DMC/DMR detection criteria were applied on P values from a beta-binomial regression model for each CpG site (the RADMeth model, column 1) or P values from PQLseq (column 2). DMCs are defined as CpG sites with Benjamini-Hochberg corrected spatially adjusted P value < 0.05, and DMRs are genomic regions with two or more consecutive Benjamini-Hochberg corrected spatially adjusted P values < 0.01 (default criteria in the current implementation of RADMeth within MethPipe version 3.4.3). Full model is the model used in the actual differential methylation analyses, including clinical and technical covariates specified in the Supplementary Table 1. The simple models were permuted sex + PC1 + PC2 and permuted epidural + sex + PC1 + PC2. These analyses were run for three permutations of each covariate.

|  | RADMeth+RADMeth | PQLseq+RADMeth |
| --- | --- | --- |
| DMCs, permuted sex, full model | 4031, 10555, 3115 | 1414, 482, 1168 |
| DMCs, permuted sex, simple model | 1352, 5747, 3870 | 1139, 598, 2390 |
| DMCs, permuted epidural, full model | 1916, 9215, 2887 | 279, 2521, 1054 |
| DMCs, permuted epidural, simple model | 559, 2910, 2466 | Not run |
| DMRs, permuted sex, full model | 189, 535, 135 | 59, 16, 50 |
| DMRs, permuted sex, simple model | 59, 297, 228 | 48, 26, 132 |
| DMRs, permuted epidural, full model | 88, 368, 139 | 8, 84, 37 |
| DMRs, permuted epidural, simple model | 21, 108, 105 | Not run |

**Table 3:**
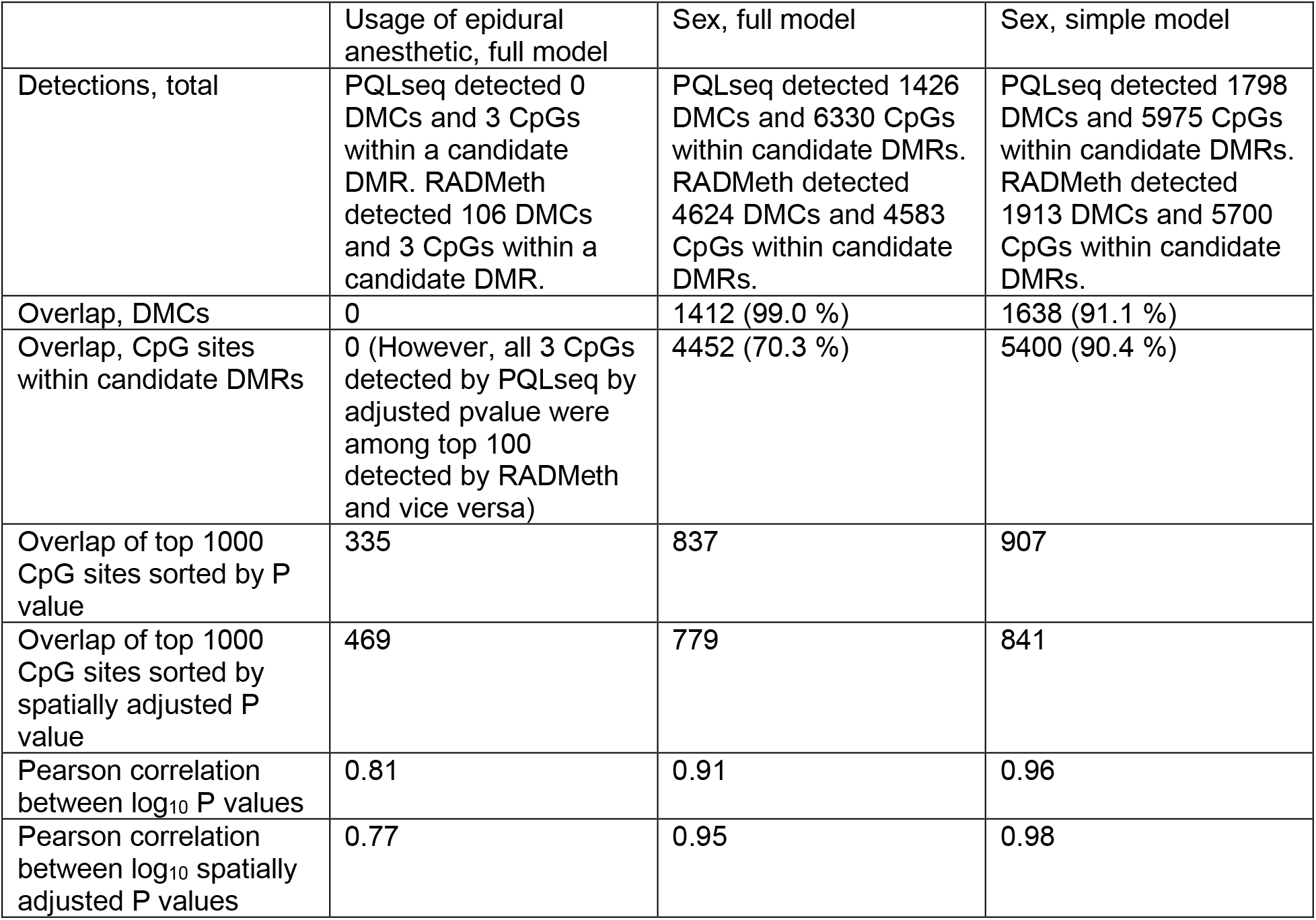
Comparison between results obtained using PQLseq (a GLMM) and RADMeth beta-binomial regression. Here, DMCs are defined as CpG sites with Benjamini-Hochberg corrected P value < 0.05 (before spatial adjustment) and CpGs within candidate DMRs are all CpG sites with empirically FDR-controlled spatially adjusted P value < 0.05. DMR detection has not been done for this comparison. The full models include all technical and clinical covariates specified in the Supplementary Table 1, and the simple model is sex + PC1 + PC2. The percentages are percentages of the detections of PQLseq.

|  | Usage of epidural anesthetic, full model | Sex, full model | Sex, simple model |
| --- | --- | --- | --- |
| Detections, total | PQLseq detected 0 DMCs and 3 CpGs within a candidate DMR. RADMeth detected 106 DMCs and 3 CpGs within a candidate DMR. | PQLseq detected 1426 DMCs and 6330 CpGs within candidate DMRs. RADMeth detected 4624 DMCs and 4583 CpGs within candidate DMRs. | PQLseq detected 1798 DMCs and 5975 CpGs within candidate DMRs. RADMeth detected 1913 DMCs and 5700 CpGs within candidate DMRs. |
| Overlap, DMCs | 0 | 1412 (99.0 %) | 1638 (91.1 %) |
| Overlap, CpG sites within candidate DMRs | 0 (However, all 3 CpGs detected by PQLseq by adjusted pvalue were among top 100 detected by RADMeth and vice versa) | 4452 (70.3 %) | 5400 (90.4 %) |
| Overlap of top 1000 CpG sites sorted by P value | 335 | 837 | 907 |
| Overlap of top 1000 CpG sites sorted by spatially adjusted P value | 469 | 779 | 841 |
| Pearson correlation between $\log_{10}$ P values | 0.81 | 0.91 | 0.96 |
| Pearson correlation between $\log_{10}$ spatially adjusted P values | 0.77 | 0.95 | 0.98 |

### Type 1 error rates were similar with different implementations of spatial adjustment and DMR detection

By default, RADMeth defines a DMR as a set of two or more consecutive CpG sites with Benjamini-Hochberg-corrected spatially adjusted P values < 0.01. With this definition, 7 – 84 DMRs were associated with each permuted covariate, when RADMeth’s spatial adjustment and DMR detection were applied to P values obtained by fitting PQLseq for the full study design (45 runs of PQLseq, each with one permuted covariate). When PQLseq was replaced by beta-binomial regression (RADMeth), these numbers were between 21 and 535 (observed for models specified in Table 2). We also tried an alternative DMR detection method, implemented as part of comb-p (11). The numbers of DMRs, that were associated with permuted usage of epidural anesthetic (random binary vectors with 50 % of the samples in each category) were between 18 and 66 (Sidák-corrected region-wise P value < 0.05), when P values from PQLseq were used as an input for comb-p, which performs spatial adjustment and DMR detection. These numbers are approximately in the same range as the corresponding numbers detected using RADMeth’s definition of a DMR (8 – 84, Table 2). All the above-mentioned spatial adjustments were done using window and step sizes recommended by RADMeth (window 200 bp, autocorrelation step size 1). Changing window size to 500 and step size to 10 had little or no effect on the results (data not shown). Thus, we conclude that different implementations of the spatial adjustment result in similar P value inflation.

### Differential methylation was reproducible between statistical models

The differential methylation models used here (the GLMM called PQLseq and the beta-binomial regression model called RADMeth) differ with respect to the noise model, optimization algorithm, and the design matrix. PQLseq includes a random effect component for the relatedness between individuals, which is not present in RADMeth. Furthermore, the design matrix needed to be modified for RADMeth such that continuous covariates were transformed to categorical (as described in Methods). Despite these differences, the results were highly concordant, as summarized in Table 3 for two example covariates: sex (which was associated with differential methylation at thousands of CpG sites) and usage of epidural anesthetic (an example of a covariate associated with very little differential methylation). For example, 85.2 % of the sex-associated DMCs that were detected by PQLseq, were confirmed by RADMeth. Since RADMeth was error-prone with the full study design, as described above, a simple model (sex + PC1 + PC2) was included in this comparison. Also, a similar comparison between results obtained using simple and full models is presented in Supplementary Table 5. The results from PQLseq were highly robust to model complexity, and results from RADMeth were robust to model complexity for the part that was empirically FDR-controlled.

### Differential methylation analysis can be improved by simple tricks that decrease missing and extreme values

The common practice to exclude values below some coverage threshold (such as 5 or 10 reads) can increase some of the above-described problems, as well as the effects of some technical biases (37). Both RADMeth and PQLseq deal with limited coverage by modeling the technical variation with a binomial distribution and therefore do not benefit from such coverage filtering. However, the quickest way to construct the input count matrices (methylated and total reads for each CpG site in each sample) from individual methylation call files of each sample, would be to first filter each methylation call file. We recommend retaining values at coverages 1 – 9, even if minimum coverage of 10 is required in a minimum number of the samples (as done here). The documentation of this workflow (40,41) includes descriptions of two simple ways to construct such a count matrix (from the output files of the Bismark methylation extractor) within a reasonable time and memory reservation, even if the number of samples is > 100. By retaining values at coverages 1 – 9, the median number of samples with missing (coverage=0) values was one (out of 173), and only 3.4 % of the CpG sites had more than 10 % missing values.

In addition to missing values, extreme values are a challenge in bisulfite data analysis. The most common methylation proportion values are 0 and 1, which are problematic in the context of generalized linear models with the logit link function (infinite in the logit-transformed space). We used a common pseudo-count transformation to avoid both extremes, as recommended for example by the developers of PQLseq (41). We tested RADMeth beta-binomial regression and PQLseq with and without the transformation. The above-described inflation of beta-binomial regression P values increased heavily in the absence of the pseudo-count transformation. For PQLseq, the pseudo-count transformation markedly improved convergence (without it the model converged at 68 – 69 % and with it at > 99 % of the 2.7 million high-coverage CpG sites), and slightly decreased the estimated type 1 error rate. The overlap of the top 1000 sex-associated CpG sites ranked by P value (between PQLseq results computed with and without the pseudo-count transformation) was 94 %.

### Sex-associated differential methylation was reproducible across studies and technologies

Differentially methylated CpG sites and regions that were obtained with the proposed permutation-based method to control FDR are summarized in Table 1 and listed in Supplementary Tables 2 and 3. A small number of differentially methylated regions/cytosines (1 – 2 DMRs and/or 1 – 10 DMCs outside the DMRs) was associated with the usage of epidural anesthetic during delivery, 1 minute Apgar points, maternal age and height, gestational weight gain, maternal smoking, and maternal insulin-treated diabetes, but not with the birth weight of the newborn infant, maternal pre-pregnancy BMI, number of earlier miscarriages, the mode of delivery, labor induction or the cosine transformed month of birth.

Altogether 1426 DMCs and 297 DMRs were associated with sex. The nearest genes were enriched in gene ontologies such as embryonic pattern specification and anatomical structure morphogenesis. All significantly enriched biological process gene ontologies (Fisher’s exact test FDR < 0.05) are listed in Supplementary Table 4. The DMR with the smallest P value (nominal P value 9.6 × 10^−24^) was on the promoter of the *PTPRF* interacting protein alpha 3 (*PPFIA3*), which interacts with the LAR family of proteins, important in mammary gland development (38). This DMR was hypomethylated in females compared to males, hence the gene is expected to be upregulated in females. Other top five differentially methylated promoter regions (ranked by P value) included the promoters of zona-pellucida binding protein 2 (*ZPBP2*) and developmental pluripotency associated 5 (*DPPA5*), which are expressed in the testis tissue and have very little or no expression in any other tissue type, according to the Genotype-Tissue Expression (GTEx) Portal on 30/11/20 (39). These regions were hypermethylated in females compared to males, hence the genes are likely to be less expressed in females. A DMR on the promoter of *ZPBP2* was selected for technical validation with targeted pyrosequencing. The hypermethylation in females of all six CpG sites was confirmed with P values in the order of 10^−6^ – 10^−9^ (Table 4). The pyrosequencing results have also been described in (10), where the sex-associated region was selected as a positive control to confirm that concordant results could be obtained with two different technologies (RRBS and targeted pyrosequencing).

**Table 4:**
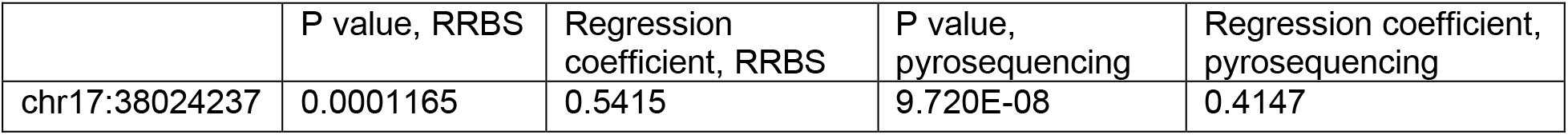
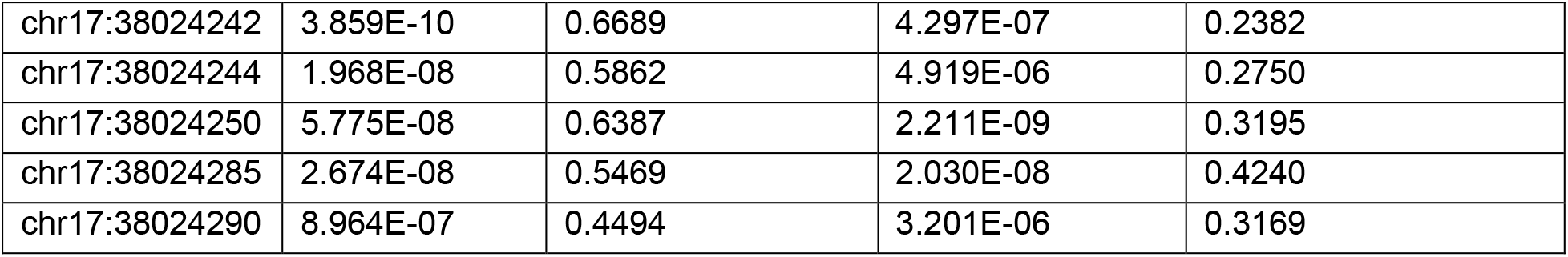
A summary of RRBS and Pyrosequencing results on the association between sex (0 = male, 1=female) and six CpG sites located on the promoter of Zona Pellucida Binding Protein 2 (ZPBP2). The RRBS data was modeled with a GLMM (PQLseq), and the pyrosequencing data was modeled with ordinary linear regression, as described in Methods.

|  | P value, RRBS | Regression coefficient, RRBS | P value, pyrosequencing | Regression coefficient, pyrosequencing |
| --- | --- | --- | --- | --- |
| chr17:38024237 | 0.0001165 | 0.5415 | 9.720E-08 | 0.4147 |
| chr17:38024242 | 3.859E-10 | 0.6689 | 4.297E-07 | 0.2382 |
| chr17:38024244 | 1.968E-08 | 0.5862 | 4.919E-06 | 0.2750 |
| chr17:38024250 | 5.775E-08 | 0.6387 | 2.211E-09 | 0.3195 |
| chr17:38024285 | 2.674E-08 | 0.5469 | 2.030E-08 | 0.4240 |
| chr17:38024290 | 8.964E-07 | 0.4494 | 3.201E-06 | 0.3169 |

We compared our results to two earlier studies on sex-associated DNA methylation in umbilical cord blood (15). Out of 390 CpG sites that were differentially methylated in our data as part of DMRs or as individual DMCs and are targeted by 450K arrays (hence could have been detected in the earlier studies), 221 CpG sites in 110 unique regions were differentially methylated according to one or both earlier studies. The overlap was highly significant (Fisher’s exact test P value < 2.2 × 10^−16^). The direction of methylation difference could not be reliably inferred from the results reported in (41) but for 192 DMCs that were common between our study and (40), the concordance of methylation difference was perfect: 154 sites were hypermethylated in females in both studies, and 36 hypermethylated in males in both studies. The high proportion of hypermethylation in females among the differentially methylated sites was highlighted by both earlier studies and further confirmed by the current study. Out of 6426 CpG sites that were either part of a DMR or differentially methylated as individual cytosines with respect to sex in our data, 73 % (or 66.2 % if each DMR is counted only once) were hypermethylated in females. Out of 221 sex-associated CpG sites that were common between our study and one or both earlier studies, six were differentially methylated as individual cytosines (Benjamini-Hochberg-corrected PQLseq P value < 0.05), 162 were differentially methylated based on the empirical FDR control of spatially adjusted P values, and 53 were differentially methylated with both criteria. CpG sites that were differentially methylated with both criteria were enriched among these confirmed findings (Fisher’s exact test P value 0.015).

## Discussion

The relatively large number of independent samples (N=173) enabled us to simultaneously model the effects of several clinical covariates and to evaluate the false discovery rate with a permutation analysis. This study might be the first one to discover the elevated type 1 error rate caused by spatial adjustment of P values with the autocorrelation-adjusted Z-test (also known as the Stouffer-Lipták-Kechris correction or comb-p (11)), implemented for bisulfite sequencing data within RADMeth (16) as part of the analysis pipeline MethPipe (15). RADMeth is a widely-used method and has performed well in simulated bisulfite sequencing data according to several independent studies (42–44). However, simulations have typically modeled only one covariate effect, which is a drastic over-simplification compared to real biomedical study settings. Furthermore, the characteristics of bisulfite sequencing data, such as large numbers of missing values and the bi-modal distribution of methylation proportions (i.e. high peaks at 0 and 1), are often not present in simulations. In earlier studies (11,16,43), real bisulfite sequencing data have been utilized to evaluate the ability of the spatial adjustment method to detect biologically meaningful differential methylation but not to evaluate its specificity. Checking the number of discoveries with permuted inputs in these data was an important sanity check that revealed the P value inflation and radically changed our conclusions on differential methylation.

The strengths of comb-p include efficiency and generalizability. It is applicable to any set of spatially correlated P values, such as those from DNA methylation microarray or bisulfite sequencing data analysis. It adjusts each P value based on neighboring P values (up to a user-defined genomic distance) and their autocorrelation, which is estimated beforehand from the whole set of P values and their genomic coordinates. Note that the original data or summary statistics, such as regression coefficients or mean methylation differences, are not used. Hence, CpG sites within DMRs (as detected by comb-p or RADMeth) are not guaranteed to have a methylation difference in the same direction. However, this can be easily checked afterwards; and for example, the DMRs reported here were filtered based on the consistency of the direction of difference. The stationarity assumption is a more important limitation. The method assumes the correlation between CpG sites at a given distance from each other to be the same across the genome. This assumption might be more realistic in DNA methylation microarray data, where most probes target promoters, than bisulfite sequencing data, which covers different types of genomic regions. Indeed, the developers of comb-p recommend a permutation analysis for type 1 error control to be performed for each unique data set that their method is applied to (11). In practice, however, it seems that this recommended sanity check is ignored in studies that apply comb-p or RADMeth for the detection of DMRs.

Besides the autocorrelation-adjusted Z-test discussed here, spatial correlation between CpG sites can be accounted for by alternative strategies; that for example, combine CpG sites to candidate regions based on the direction of difference between two groups before the actual differential methylation analysis (45) or directly model the autocorrelation structure between CpG-sites as a random effect within each input region (44). The reason why we chose a GLMM (PQLseq) that was fit separately for each CpG site (40,41), followed by a simple Stouffer-Lipták-Kechris correction for the spatial adjustment, was computational efficiency compared to other approaches and the ability to include both binary and continuous covariate effects.

Appropriate multiple testing correction is vital in bisulfite sequencing based DNA methylation studies, which typically include millions of CpG sites. Methods such as RADMeth beta-binomial regression and PQLseq fit a model separately for each CpG site. Performing a FWER correction (such as Bonferroni) or an FDR correction (Benjamini-Hochberg) on the P values of individual CpG sites would be overly conservative and ignore the spatial correlation, which is often high, up to a distance of approximately 1000 base pairs (46). Furthermore, DMRs are biologically more interpretable than individual DMCs. The spatial adjustment of individual P values is therefore useful, but the type 1 error control needs careful consideration. The empirical approach taken in this study is an efficient guard against any type of P value inflation. We recommend a similar permutation analysis for future studies, whenever the number of samples is large enough. Another option is to determine the significance based on P values that have not been spatially adjusted; applying for example, the epigenome-wide significance threshold (47) or the Benjamini-Hochberg correction to detect individual DMCs. If Benjamini-Hochberg (or some equivalent approach) is chosen, the significance threshold might be slightly relaxed to compensate for the fact that the true number of independent tests is much smaller than the total number of CpG sites (which is typically 2 – 3 million in RRBS data). Efficient implementations for DMR detection, such as comb-p or RADMeth, can then be used to define regions around the DMCs.

The empirical FDR control strategy has some obvious limitations. A proper permutation test would require repeating the time-consuming and memory-intensive differential methylation analysis and spatial adjustment for 2.7 million CpG sites at least 1000 times for each covariate of interest. Since this is not feasible, we do not report false discovery rates for each CpG site. Instead, a cutoff is chosen such that the false discovery rate is estimated not to exceed 0.05.

Here the estimates are based on three permutations of each covariate. Keeping this limitation in mind, each permuted vector was created such that it did not correlate with any actual clinical or technical variable. This way the permuted study designs mimicked the design matrix used in our analysis (which did not include strongly correlated covariates) and the results were likely to represent relevant null distributions. Results must be interpreted cautiously, especially if less than 20 CpG sites are below the empirically determined threshold, in which case the empirically estimated FDR would be zero – a definite sign of a too low number of permutations (48).

## Conclusions

Here, we have developed a bisulfite sequencing data analysis workflow, which is available (18) and completely based on free open source tools. We have shown that Stouffer-Lipták-Kechris corrected P values are inflated, and both implementations of DMR detection (comb-p and RADMeth) find false positives in bisulfite sequencing data, if default significance thresholds are applied. Based on empirically estimated thresholds, very little differential methylation was associated with any of the included variables, other than sex. The results on sex-associated DNA methylation were highly reproducible across different analysis workflow strategies, as well as independent studies. A large proportion of perinatal sex-associated epigenetic differences reported in earlier studies (40,41) were confirmed by this study. With the RRBS method, we were also able to report thousands of novel sex-associated differentially methylated cytosines that could not have been detected in the earlier studies, which were limited to cytosines included in the Illumina 450K DNA methylation microarrays. Technical validation of a novel sex-associated promoter region by targeted pyrosequencing further supported the validity of these results.

## List of abbreviations

CpG: A genomic site where cytosine (C) is followed by guanine (G). The p stands for phosphate which connects two adjacent bases in the genome.
DMC: Differentially methylated cytosine
DMR: Differentially methylated region
FDR: False discovery rate
GLMM: Generalized linear mixed effects model
RRBS: Reduced representation bisulfite sequencing
PC1 and PC2: Projections of the sample-specific methylation proportion vectors on the first two orthonormal principal components of the methylation proportion matrix

## Declarations

### Availability of data and materials

The dataset supporting the conclusions of this article will be available in ArrayExpress, accession code E-MTAB-10530, as soon as the related preprint (10) is accepted for publication. Pre-processed RRBS data (numbers of methylated and total numbers of reads at each high-coverage CpG site in each sample), as well as the data on technical variables, sex, and the disease outcome (development of multiple type 1 diabetes associated autoantibodies before age 15) will be available. To protect the privacy of the study participants, data on the other variables are only accessible from the corresponding author of (10) upon reasonable request. The documentation of the data analysis workflow, mostly written in R 4.0.4, is available in GitHub (18), where also the data availability status will be updated.

### Author contributions

E.L. developed the analysis workflow, analyzed the data, interpreted the results, prepared the figures and tables, and wrote the manuscript. V.H participated in the development of the analysis workflow and the interpretation of the results T.G. was responsible for technical validation by targeted pyrosequencing. U.U. participated in the interpretation of the results. M.N., M.V-M., and J.M. provided the clinical information compiled by E.L., H.K., and N.L.. O.R. supervised the laboratory experiments. J.T. provided the clinical information and supervised M.N. and M.V-M.. M.Kn and M.O. initiated the study together with R.La.. R.Lu. was responsible for the bisulfite sequencing and participated in the interpretation of the results. R.La. supervised the study and participated in the interpretation of the results, H.L. supervised E.L. and V.H., participated in the interpretation of the results, and revised the manuscript. All authors contributed to the final version of the manuscript.

## Acknowledgements

We are grateful to the personnel of Turku University Hospital. We thank Riitta Veijola, Jorma Ilonen, and Heikki Hyöty for providing the data from the Diabetes Prediction and Prevention (DIPP) study. We thank Mikko Konki and Roosa Kattelus for assistance in the Pyrosequencing. We are grateful to Bishwa R. Ghimire, Asta Laiho, and Laura L. Elo for their insight into the RRBS data analysis. We acknowledge the Turku Bioscience Centre’s core facility, the Finnish Functional Genomics Centre (FFGC) supported by Biocenter Finland, for their assistance. We acknowledge the Finnish Centre for Scientific Computing (CSC) and the computational resources provided by the Aalto Science-IT project.

