## Supplementary Table 1 for "Permutation-based significance analysis reduces the type 1 error rate in bisulfite sequencing data analysis of human umbilical cord blood samples"

Supplementary Table 1: Clinical and technical covariates

| Covariate(s) | Included in regression (yes/no) | Details and reasons for inclusion/exclusion |
|---|---|---|
| Age, mother | Yes | Maternal age only correlates with the number of earlier pregnancies and deliveries, and is prioritized over them, since continuous covariates are easier to model than counts. |
| Birth length | No | Correlates with birth weight |
| Birth weight | Yes | Birth weight is included to represent the size of the newborn infant. It is prioritized over birth height, head circumference and pregnancy duration, since birth weight has the best measurement accuracy. |
| BMI, mother | Yes | This is the pre-pregnancy body mass index. It is prioritized over the body weight, since BMI is more comparable between individuals |
| Caesarean section | Yes | The mode of delivery was simplified to vaginal/C-section. C-section only correlates with perinatal asphyxia and is prioritized, since it is simple to define and does not include any measurement uncertainty |
| Duration of delivery stage 1 | No | Difficult to measure and includes too many missing values (32 out of 173), for example at all C-section cases. Also correlates with the usage of epidural anesthetic |
| Duration of delivery stage 2 | No | Too many missing values (28 out of 173), for example at all C-section cases. Also correlates with the usage of epidural anesthetic |
| Gestational vaginal bleeding | No | A binary variable, correlates with induced labor |
| Gestational weight gain, mother | Yes | Does not correlate with any other covariate |
| Glucose tolerance test result, mother | No | A binary variable, correlates with insulin-treated diabetes and includes many missing values (137 out of 173) |
| Head circumference | No | Correlates with birth weight |
| Height, mother | Yes | Included to represent the maternal size together with the pre-pregnancy BMI. Maternal height correlates only with maternal weight. |
| Induced labor | Yes | Correlates only with gestational vaginal bleeding and is prioritized, since induced labor is simple to define, whereas gestational vaginal bleeding can have different degrees of severity |
| Insulin-treated diabetes, mother | Yes | This can be insulin-treated diabetes of any type (for example gestational diabetes). This covariate is prioritized over neonatal intensive care, neonatal hypoglycemia, and earlier C-section, since insulin-treatment throughout the pregnancy is expected to be more relevant for umbilical cord blood than events that take place before or after the pregnancy. |
| Islet cell autoantibody positivity before age 15 ("Class") | Yes | Differences between newborn infants who later progress to type 1 diabetes and controls who remain autoantibody-negative are reported in the preprint by Laajala et al. (1) for 122 individuals, who are a subset of the 173 individuals in this study. This study includes 51 individuals, who were excluded from the study on type 1 diabetes due to transient autoantibody positivity or positivity for only one biochemical autoantibody. For this study, islet autoantibody positivity was simplified to yes/no (1=persistent positivity for one or more |

| | | |
|---|---|---|
| | | biochemical autoantibody, 0=no persistent autoantibody positivity). |
| Library preparation batch | Yes | This is a categorical variable with 7 levels (transformed to 6 binary covariates + intercept). A median number of 23 samples (range 5 – 48) were processed within each batch. |
| Low Apgar points | Yes | The 1-minute Apgar points were used here as a binary variable (0=normal, 1=low). Values 7 and lower were considered low. |
| Multiple pregnancy | No | This binary variable correlates with birth weight |
| Neonatal hypoglycemia | No | Correlates with maternal insulin-treated diabetes |
| Neonatal intensive care | No | Correlates with maternal insulin-treated diabetes |
| Number of earlier C-sections | No | The number of earlier Caesarean sections was simplified to a binary variable (0=none, 1=one or more). It correlates with maternal insulin-treated diabetes. |
| Number of earlier deliveries | No | The number of earlier deliveries was simplified to a binary variable (0=none, 1=one or more). It correlates with maternal age. |
| Number of earlier miscarriages | Yes | The number of earlier miscarriages was simplified to a binary variable (0=none, 1=one or more). There was no reason to exclude this covariate, since it only correlated with the number of earlier pregnancies. |
| Number of earlier pregnancies | No | The number of earlier pregnancies was simplified to a binary variable (0=none, 1=one or more). It correlates with maternal age. |
| Perinatal asphyxia | No | Correlates with Caesarean section |
| Pregnancy duration | No | Correlates with birth weight |
| Principal components 1 and 2 | Yes | Principal components 1 and 2 were included in the model to represent technical variation. |
| Sex | Yes | Binary, 0=male, 1=female. Correlates with no other covariate |
| Smoking during pregnancy | Yes | Covariates related to in-utero conditions were generally prioritized. |
| Transformed month | Yes | This is the month of birth, cosine-transformed to season of birth ($\cos(2\pi m/12)$, where m is the month as numbers 1 – 12) |
| Umbilical arterial blood pH | No | Too many missing values (23 out of 173) |
| Usage of epidural anesthetic | Yes | This is a binary variable indicating whether epidural anesthetic was used during delivery stage 1. This variable was prioritized over duration of delivery stages 1 and 2 since usage of epidural anesthetic is simple to define and does not include missing values or measurement uncertainty. |
| Weight of the newborn infant when leaving the hospital | No | Correlates with birth weight and is not easily comparable between different individuals, since they spend variable amounts of time in the hospital |
| Weight, mother (initial and final) | No | Correlates with maternal pre-pregnancy BMI and height. Initial weight is the pre-pregnancy weight and final weight is the last recorded weight before delivery. |
| Year of birth (= sample collection year) | Yes | Included to account for possible technical variation related to storage time in -20°C |

1. Laajala E, Ullah U, Grönroos T, Rasool O, Halla-aho V, Konki M, et al. Umbilical cord blood DNA methylation in children who later develop type 1 diabetes. medRxiv. 2021.
