## Supplementary Table 5 for "Permutation-based significance analysis reduces the type 1 error rate in bisulfite sequencing data analysis of human umbilical cord blood samples"

Supplementary Table 5: Comparison between results obtained by fitting the full vs. the simple model. The full models included all clinical and technical covariates specified in Supplementary Table 1. The simple models were sex + PC1 + PC2 (columns 1 and 2, testing for sex) and epidural + sex + PC1 + PC2 (column 3, testing for epidural). Here, DMCs are defined as CpG sites with Benjamini-Hochberg corrected P value < 0.05 (before spatial adjustment) and CpGs within candidate DMRs are all CpG sites with empirically FDR-controlled spatially adjusted P value < 0.05. DMR detection has not been done for this comparison. The percentages are percentages of the detections with the full model.

| | PQLseq, sex | RADMeth, sex | RADMeth, usage of epidural anesthetic |
|---|---|---|---|
| Detections, total | Altogether 1426 DMCs and 6330 CpG sites within candidate DMRs were detected by fitting the full model. Fitting the simple model, 1798 DMCs and 5975 CpG sites within candidate DMRs were detected. | Altogether 4624 DMCs and 4583 CpG sites within candidate DMRs were detected by fitting the full model. Fitting the simple model, 1913 DMCs and 5700 CpG sites within candidate DMRs were detected. | Altogether 106 DMCs and 3 CpG sites within a candidate DMR were detected by fitting the full model. Fitting the simple model, 3 DMCs and 0 CpG sites within candidate DMRs were detected. |
| Overlap, DMCs | 1215 (85.2 %) | 1841 (39.8 %) | 0 |
| Overlap, CpG sites within candidate DMRs | 5476 (86.5 %) | 4466 (97.4 %) | 0 |
| Overlap of top 1000 CpG sites sorted by P value | 810 | 797 | 285 |
| Overlap of top 1000 CpG sites sorted by spatially adjusted P value | 861 | 832 | 392 |
| Pearson correlation between $\log_{10}$ P values | 0.88 | 0.88 | 0.77 |
| Pearson correlation between $\log_{10}$ spatially adjusted P values | 0.95 | 0.94 | 0.75 |
